# Coral environmental history is primary driver of algal symbiont composition, despite a mass bleaching event

**DOI:** 10.1101/2022.05.24.493152

**Authors:** Mariana Rocha de Souza, Carlo Caruso, Lupita Ruiz-Jones, Crawford Drury, Ruth D. Gates, Robert J. Toonen

## Abstract

Coral reefs are iconic examples of climate change impacts because climate-induced heat stress causes the breakdown of the coral-algal symbiosis leading to a spectacular loss of color, termed ‘coral bleaching’. To examine the fine-scale dynamics of this process, we re-sampled 600 individually marked *Montipora capitata* colonies from across Kāne‘ohe Bay, Hawai’i and compared the algal symbiont composition before and after the 2019 bleaching event. The relative proportion of the heat-tolerant symbiont *Durusdinium* in corals increased in most parts of the bay following the bleaching event. Despite this widespread increase in abundance of *Durusdinium*, the overall algal symbiont community composition was largely unchanged, and hydrodynamically defined regions of the bay retained their distinct pre-bleaching compositions. Furthermore, depth and temperature variability were the most significant drivers of Symbiodiniaceae community composition by site regardless of bleaching intensity or change in relative proportion of *Durusdinium*. Our results suggest that the plasticity of symbiont composition in corals may be constrained to adaptively match the long-term environmental conditions surrounding the holobiont, independent of an individual coral’s stress and bleaching response.

## 1. Introduction

Anthropogenic climate change impacts ecosystems across the globe (1). Coral reefs are among the most iconic examples of climate-driven ecosystem decline, exhibiting a characteristic loss of color called ‘bleaching’ in response to thermal stress. Bleaching is the paling of corals resulting from the breakdown of the symbiosis between the cnidarian host and dinoflagellate algae of the family Symbiodiniaceae, which is responsible for meeting about 90% of reef-building coral energy requirement (2). Because corals are metabolically dependent on this symbiosis, long periods in a bleaching state can deplete host energy supply and reserves (3–5), impact coral growth (6, 7) reproduction (8–10), and result in coral mortality (11–13). The frequency of mass coral bleaching events worldwide has increased nearly fivefold in the past four decades (14–17), resulting in significant losses of live coral in many parts of the world (18, 19). Despite visual recovery, the impacts of bleaching may persist for years (10, 20) and the increasing frequency and duration of marine heatwaves suggests there might not be enough time for corals to recover between bleaching events (18, 21). Ocean temperatures are predicted to rise 1-2 ^o^C under best-case emission scenarios (22), and coral persistence through increasingly frequent and severe heatwaves is dependent on the capacity to acclimatize or adapt to a rapidly changing environment (14, 23–25). These factors make coral reefs one of the most vulnerable ecosystems to increasing global temperatures (26, 27).

One mechanism by which corals may deal with thermal stress is through a change in the relative proportion of more thermally resistent algal endosymbionts hosted by corals that experience thermal stress (21, 28). Although some corals maintain stable associations or revert to pre-bleaching algal symbiont composition (29, 30), others change their algal symbiont community composition following bleaching events and can maintain altered proportions of algal symbionts after recovery from bleaching (31, 32). The association of coral with specific types of Symbiodiniaceae can directly influence how corals respond to environmental stress (33–36). For example, corals dominated by Symbiodiniaceae from the genus *Durusdinium* (previously clade D; 37) tend to be more resilient to heat stress and thus experience less bleaching (38–40), including in our study species, *Montipora capitata* (41, 42). Despite increasing resistance to bleaching, hosting the stress tolerant *Durusdinium* often comes at an energetic cost, decreasing the growth or metabolite exchange rate of the host, although a suite of both biotic and abiotic factors can modify such generalizations (41).

While differential susceptibility to bleaching mortality among coral species is relatively well documented (12, 43–45), there is also ubiquitous intraspecific variation in coral bleaching (46–53). Whereas temperature and irradiance are generally accepted to be the main environmental factors contributing to coral bleaching severity (18, 54–57), bleaching severity is often highly variable among individuals within and among nearby sites (41, 44, 58–60). Bleaching severity can also be influenced by the environmental conditions the corals experienced before or during the heat stress (47, 61–65). Further, other environmental factors that can modify bleaching responses of corals often correlate with irradiance and temperature, such as depth (60, 66), sedimentation (67, 68), wave energy (69), and flow (70). Other factors such as suspended sediments (71, 72), nutrient input (73, 74), and acidification (75, 76) can exacerbate or ameliorate coral bleaching severity. Because these factors contribute to the breakdown of the coral-algal symbiosis, it seems likely that they may also impact the community composition of algal symbionts, but there is no strong consensus about the role of environmental drivers of spatial variation in coral algal symbiont community structure (77–81).

Kāne‘ohe Bay, the largest protected embayment in the Hawaiian Islands, has among the highest coral cover in Hawai‘i (82, 83). The bay is environmentally and biologically heterogeneous (84–86), with the northern extents experiencing higher circulation and lower mean residence times than the southern portion of the bay (87, 88). Interestingly, because Kāne‘ohe Bay is relatively shallow and has high productivity and long residence times, the fluctuations in pCO_2_ and temperature are increased relative to open coastal reefs. corals in Kāne‘ohe Bay are are exposed to temperature and acidification regimes that will not be seen for decades in other parts of the state (84), leading to divergent environmental tolerances between the corals growing in Kāne‘ohe Bay and conspecifics collected from exposed coastal reefs a few kilometers away (89). Further, thermal tolerance experiments conducted in 2017 show that individuals take longer to bleach, maintain higher calcification rates, and experience lower bleaching mortality, than were observed for the same species at the same location in 1970 (24). Cumulatively, these results suggest corals in Kāne‘ohe Bay have become more resistant to thermal stress, and may indicate an important role of environmental history in improving stress tolerance and susceptibility to coral bleaching.

The rice coral, *Montipora capitata*, is a dominant reef builder in Kāne‘ohe Bay, where it hosts symbiodinium in the genera *Cladocopium, Durusdinium* or a mixed community (41, 42, 90–92). The environmental heterogeneity of the bay and the complexity of symbiont communities in *M. capitata*, create an ideal system to investigate factors influencing coral bleaching response and resilience (84, 89, 93, 94). We previously quantified the algal symbiont across an environmental mosaic throughout Kāne‘ohe Bay in 2018 (81) before using this baseline to re-sample colonies after the 2019 bleaching event (95) to compare algal symbiont community change through time. Here, we take advantage of this natural bleaching event to examine whether Symbiodiniaceae community structure changes in response to thermal stress, and if so, whether environmental factors modify the community response of algal symbionts within individual *M. capitata* colonies across the environmental mosaic of Kāne‘ohe Bay.

## 2. Material and Methods

### 2.1 SITE SELECTION AND TAGGING

*Montipora capitata* colonies were tagged on 30 patch reefs in Kāne‘ohe Bay, O’ahu, Hawai‘i, under SAP permit 2018-03 and SAP 2019-16 to HIMB from Hawai‘i Department of Aquatic Resources. Kāne‘ohe Bay was divided into 5 ‘blocks’ based on modeled water flow regimes and water residence times (88) and six sites (86) were selected in each block using stratified random sampling within habitats designated as patch reefs (**Figure 1**). Site IDs consist of the digit corresponding to the block in which the site is contained, followed by the site number (e.g., 1_10, with six sites per block, but not necessarily in consecutive order).

**Figure 1.**
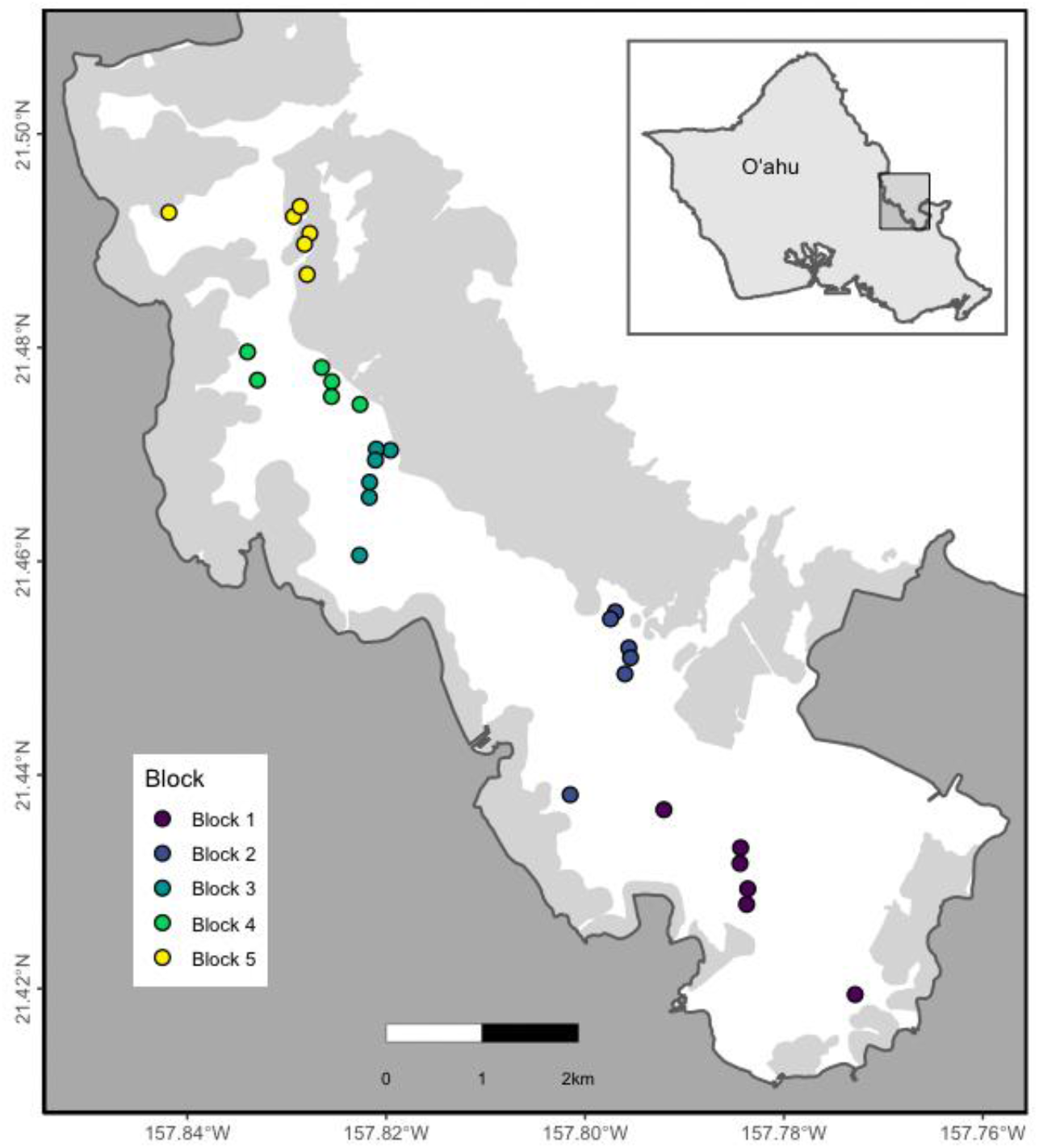
Map of Kāne‘ohe Bay highlighting the location of each of the 30 sites in the bay.

Temperature loggers (Hobo Pendant from Onset Computer Corp: UA-001-64 Data Logger) were deployed at the center of each site. Temperature recordings every 10 min began on 12 July 2017 and continued until 26 July 2019, with the loggers periodically retrieved and recalibrated throughout the study period. Sediment traps were also deployed at the center of the block and exchanged every 1-2 months, and the weight of sediment was used to estimate the sedimentation rate in each site following (96). See (86) and (81) for details on site selection, host genetic sampling, and environmental data collection and analysis.

Twenty *M. capitata* colonies were tagged at each site and 1cm^2^ clippings of each colony were collected in 2018 from visually healthy colonies. During the 2019 bleaching event, the colonies were re-visited between 3 and 21 October 2019, photographed and recollected. Sampled fragments were immediately preserved in 70% ethanol and stored at −20°C until processed. DNA from colonies collected in both years was extracted using the Nucleospin Tissue Kits (Macherey-Nagel, Düren, Germany) following manufacturer’s instructions.

During field sampling in October 2019, corals were assigned a visual bleaching score (24, 52, 93, 97): (0) totally bleached (>80% of colony white with no visible pigmentation); (1) pale (60 - 80% colony affected by pigment loss); (2) pale (10-50% colony affected by pigment loss); (3) fully pigmented (<10% colony with any visible paling) (52). Each colony was scored two times independently by two different observers using *in situ* photographs taken during collection and the mean value was assigned as the bleaching score.

### 2.2 SYMBIODINIACEAE ITS2 AMPLICON SEQUENCING LIBRARY PREPARATION

ITS2 amplicon libraries were prepared from extracted DNA and sequenced following de Souza et al. (98). Briefly, the ITS2 region was amplified for each sample, pooled and sequenced on an Illumina MiSeq platform (v3 2×300bp PE) at University of Hawaii at Manoa. Raw sequences were demultiplexed and quality filtered using Cutadapt (99). To ensure differences in read number did not impact results or interpretation, we excluded 20 samples from 2018 and 24 from 2019 whose number of reads were more than 2 standard deviations above or below the mean. Forward and reverse reads were submitted to SymPortal (100), a platform for identifying Symbiodiniaceae using high throughput ITS2 sequence data that differentiates intra- and intergenomic sources of ITS2 sequence variance. Sets of ITS2 sequences, occurring in a sufficient number of samples within both the dataset being analyzed and the entire database of samples run through SymPortal were identified as ‘defining intragenomic variants’ (DIVs) which were then used to characterize ITS2 type profiles.

In this study, we analyzed data based on Symportal outputs for Symbiodiniaceae “type” and “profile”. A type refers to Symbiodiniaceae taxa that have a specific sequence as their most abundant sequence. A Symbiodiniaceae profile is a summary description set of ITS2 sequences that have been found co-occurring in a sufficient number of samples (DIV).

### 2.3 STATISTICAL ANALYSIS

All analyses were completed in R 2021.09.0+351 version (R Core Team, 2020). We calculated a variety of summary statistics from the temperature time series for each site (86): mean daily temperature, average daily range, mean daily standard deviation, global mean, maximum and minimum temperature at each site. We calculated Degree heating weeks (DHW) per site as the accumulated time when temperature was above the bleaching threshold, set as 28.5°C given a MMM of 27.5 ^o^C (42, 101).

To examine if the relative proportion of heat resistant algal symbionts changed following a bleaching event, we calculated the relative proportion of *Durusdinium* in each colony from relative abundance values in 2018 and 2019 (81) and analyzed these data by site and block.

We used non-metric multidimensional scaling (NMDS) and permutational analysis of variance (PERMANOVA) in the R package *vegan* to examine symbiont community differentiation (Bray-Curtis dissimilarities) by year, block and site nested within block. We used the function pairwise.adonis to compare each block in 2018 and in 2019 and visualized the R^2^ from the PERMANOVA in a dendrogram using the package *pheatmap*.

To examine the influence of environmental factors on Symbiodiniaceae composition, we used a distance-based redundancy analysis (dbRDA) based on Bray-Curtis distances and tested the significance of each environmental driver using *vegan*. We calculated variance explained by each environmental variable as a proportion of total variance explained by environment (i.e., excluding residuals). We designated colonies with a relative abundance of >80% from a single genus as majority *Cladocopium* (C) or *Durusdinium* (D), with all remaining samples designated as mixed CD, corresponding to corals with no dominant algal symbiont genus.

## 3. Results

### 3.1 SYMBIODINIACEAE IDENTIFICATION

In 2019, 496 colonies passed initial quality control steps, representing a loss of ~100 colonies due to missing tags, mortality, or other unknown causes. The relative representation of types, profiles and symbiont genera were broadly similar across years, representing the background distribution of Kāne‘ohe Bay. A total of 214 Symbiodiniaceae types were identified in 2019, 178 (83%) in the genus *Cladocopium* and 36 (17%) in the genus *Durusdinium*. These numbers are similar to results from 2018, where 283 Symbiodiniaceae types were identified, 241 (85%) belonging to *Cladocopium*, and 42 (15%) belonging to the *Durusdinium* (81). Twenty-nine ITS2 DIV profiles were identified across all samples in 2019, with twenty-five belonging to the genus *Cladocopium* and 4 belonging to the genus *Durusdinium*, consistent with 2018 when twenty-six ITS2 type profiles were identified across all samples, twenty-three of which were from the genus *Cladocopium*, with the remaining 3 belonging to the genus *Durusdinium*.

In 2019, 30% of colonies hosted *Durusdinium* only, an increase of 19% from 2018 when 11% hosted only *Durusdinium*. In 2019, 22% of colonies hosted a mixed community of both genera, 24% fewer than 2018, when 46% of colonies hosted a combination of both genera. In 2019, among mixed colonies (N=109), 22 (20%) were dominated by *Cladocopium* (>80% of reads identified as *Cladocopium)*, and 38 (35%) were dominated by *Durusdinium* (>80% of reads identified as *Durusdinium)*, while the remaining 49 (45%) had moderate abundances of both genera.

### 3.2 SYMBIODINIACEAE COMMUNITY COMPOSITION BEFORE AND AFTER BLEACHING

Corals at most sites had a combination of *Cladocopium* and *Durusdinium* regardless of year (**Figure 2,** Supplemental material **Figure 1**). Sites 5_3 and 5_6 in block 5 (the most northern part of the bay) were exceptions where we did not find *Durusdinium* in either year. In 2019, there was a general increase in the proportion of *Durusdinium* in sites located in all blocks following the bleaching event (**Figure 2, 3A**); however, there was considerable variation between sites. For example, only site 5_1 in block 5 had an increased proportion of *Durusdinium*, while the other sites in block 5 remained nearly constant through time. Similarly, there was substantial variation in the change in proportion of *Durusdinium* between from 2018 to 2019 in each block (**Figure 3B,** Supplemental material **Figure 2A, B)**, with block 4 presenting the highest increase, while block 2 and 5 had limited or no increases in *Durusdinium*. In each year, the proportion of *Durusdinium* hosted by colonies was lower in the northern and southern extremes of the bay (**Figure 2**), which have relatively unique environments.

**Figure 2.**
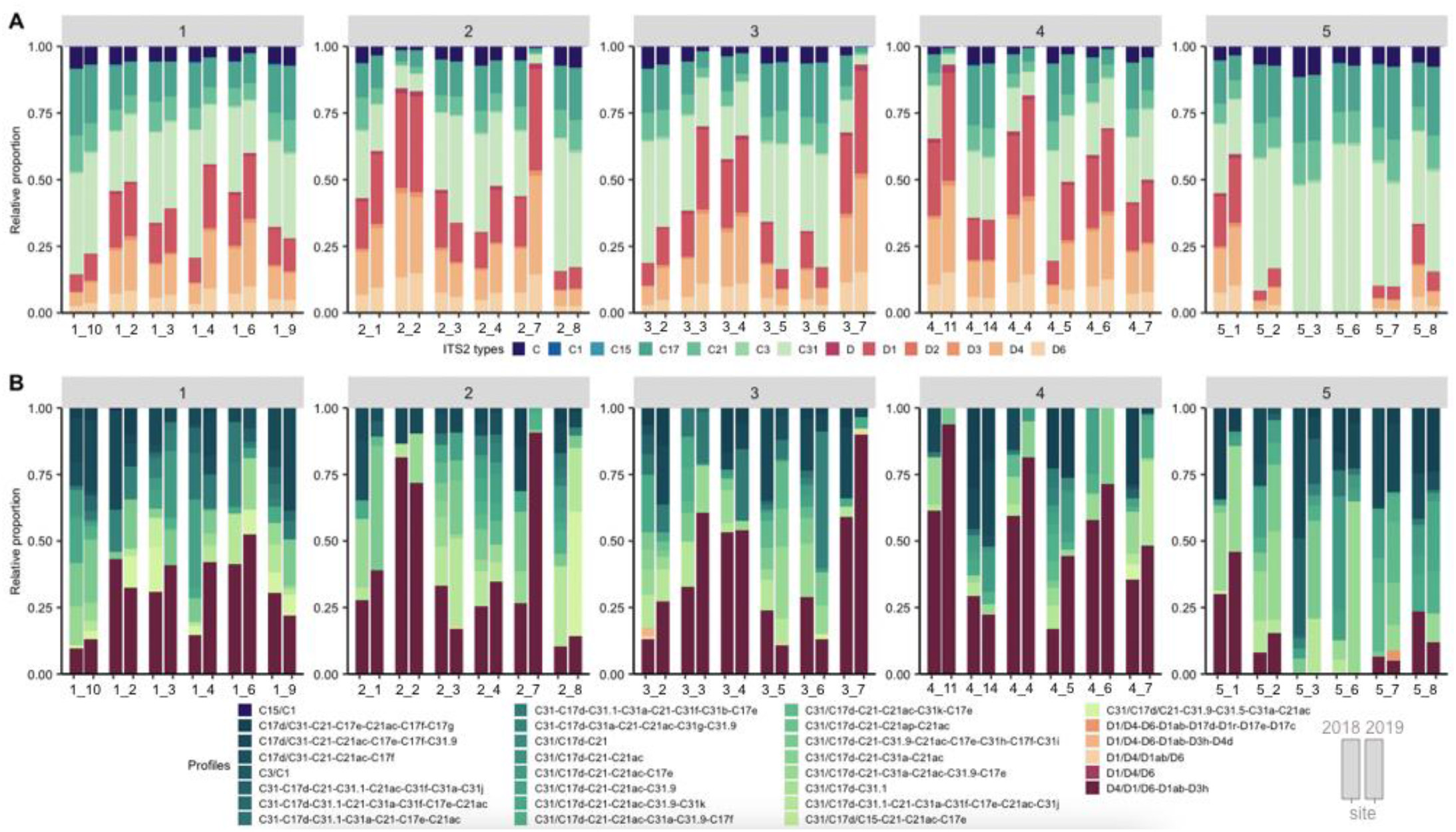
*Montipora capitata* Symbiodiniaceae community composition found in each of the 30 sites in Kāne‘ohe Bay for A) types and B) profiles. A type refers to Symbiodiniaceae taxa that have a specific sequence as their most abundant sequence. A Symbiodiniaceae profile is a summary description set of ITS2 sequences that have been found in a sufficient number of samples (DIV). Here pairs of bars per site shows the algal symbiont community in 2018 and 2019, respectively. Symbiodiniaceae ITS2 subtypes were summarized to the major subtype to facilitate visualization in the bar charts (i.e., C31a and C31b were summarized as C31). Due to the wide diversity of ITS2 available in the SymPortal database, not all sequences are given names. Only DIV sequences are named, so unnamed *Cladocopium* and *Durusdinium* sequences were combined for visualization and represented as summed “C” and “D” types, respectively.

**Figure 3.**
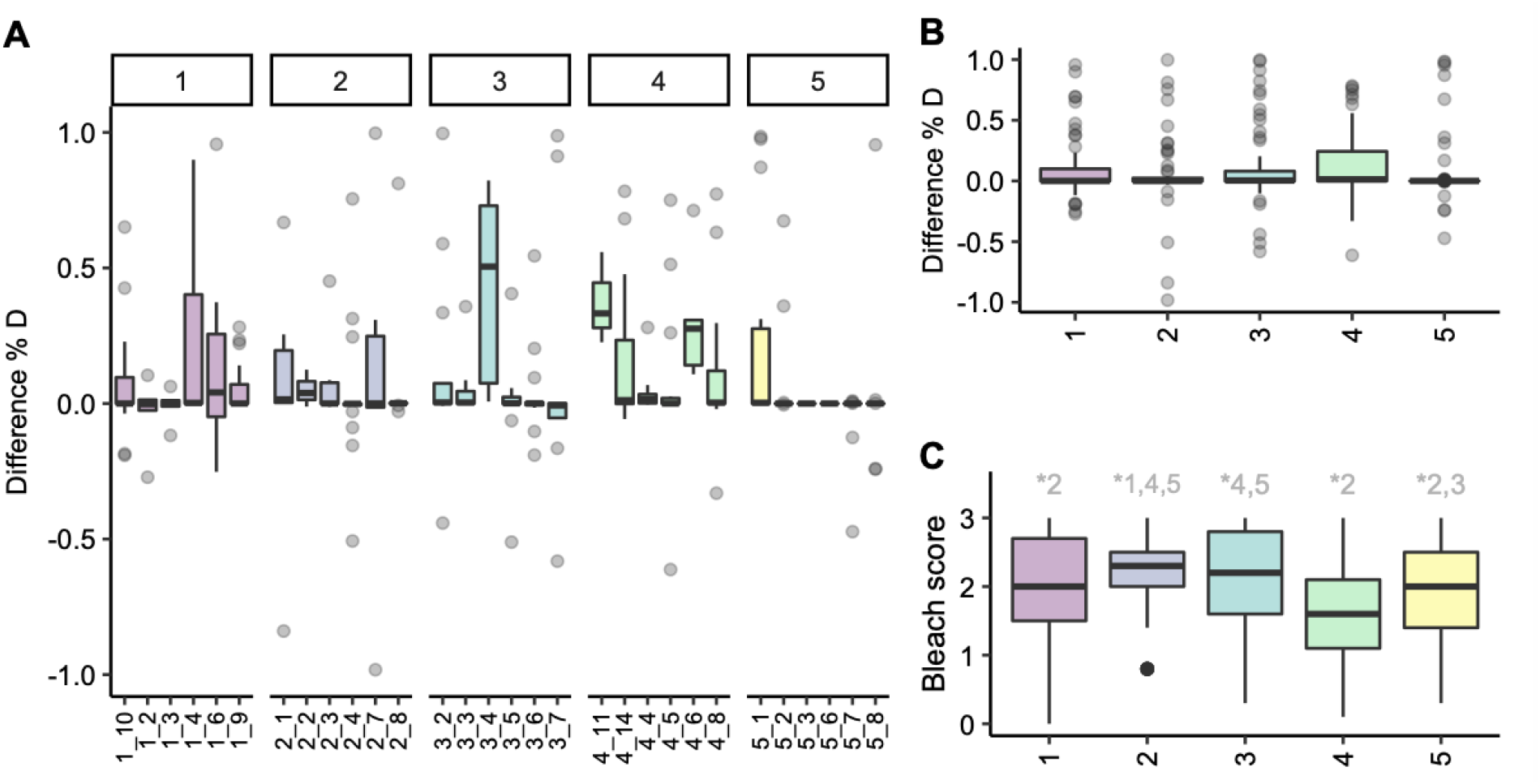
Change in *Durusdinium* in *M. capitata* in Kāne‘ohe Bay in 2018 and 2019. A) Boxplot of difference of proportion of *Durusdinium* (D) in 2019 and 2018 in each site, in each of the 5 blocks. B) Boxplot of difference of proportion of *Durusdinium* (D) in 2019 and 2018 in each of the 5 blocks. C)Bleach score per block (3: healthy, 0: fully bleached); numbers at the top of panel C represent significant comparisons from pairwise PERMANOVA.

Consistent with many other studies, bleaching stress resulted in a relative increase of *Durusdinium* among colonies of *M. capitata* across Kāne‘ohe Bay in 2019 (**Figure 3A,B**). Bleaching score (**Figure 3C,** Supplemental material **Figure 2C**) was significantly different between blocks, with the highest average score in block 4, and and lowest average score in block 2. Bleaching severity was negatively related to proportion of *Durusdinium* (Supplemental Material **Figure 3**; p<0.01). Mean bleaching score in October 2019 for the 30 sites we surveyed in the bay was 2.2 (corresponding to paling), suggesting that *M. capitata* in the 2019 bleaching event experienced milder consequences compared to bay-wide estimates for previous bleaching events in which 62% (1996), 45% (2014) and 30% (2015) of colonies bleached (82). Overall algal symbiont composition was significantly different between years (PERMANOVA F _9_= 16.322, *p*= 0.001; **Figure 4C**, **Table1**).

**Figure 4.**
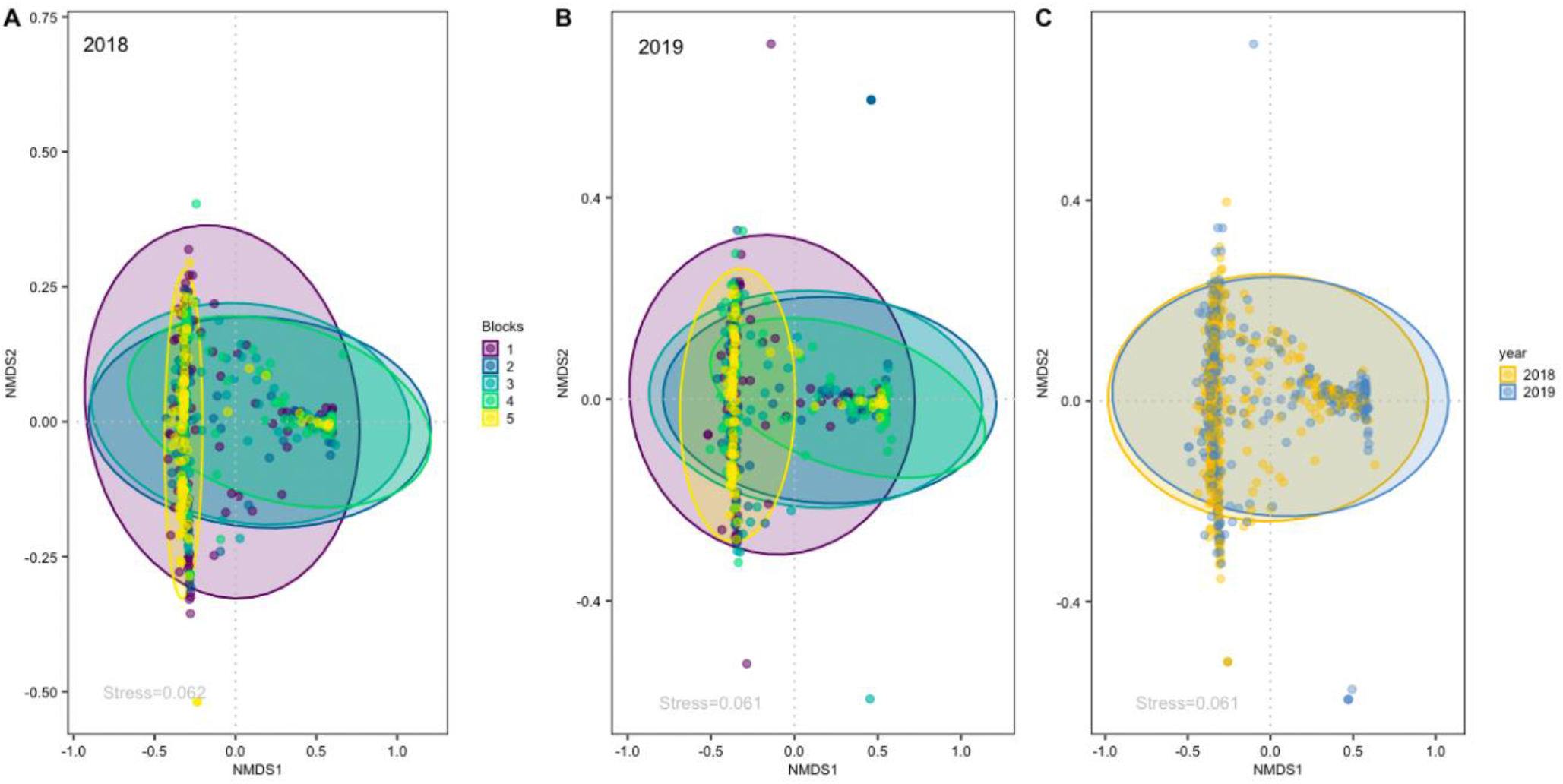
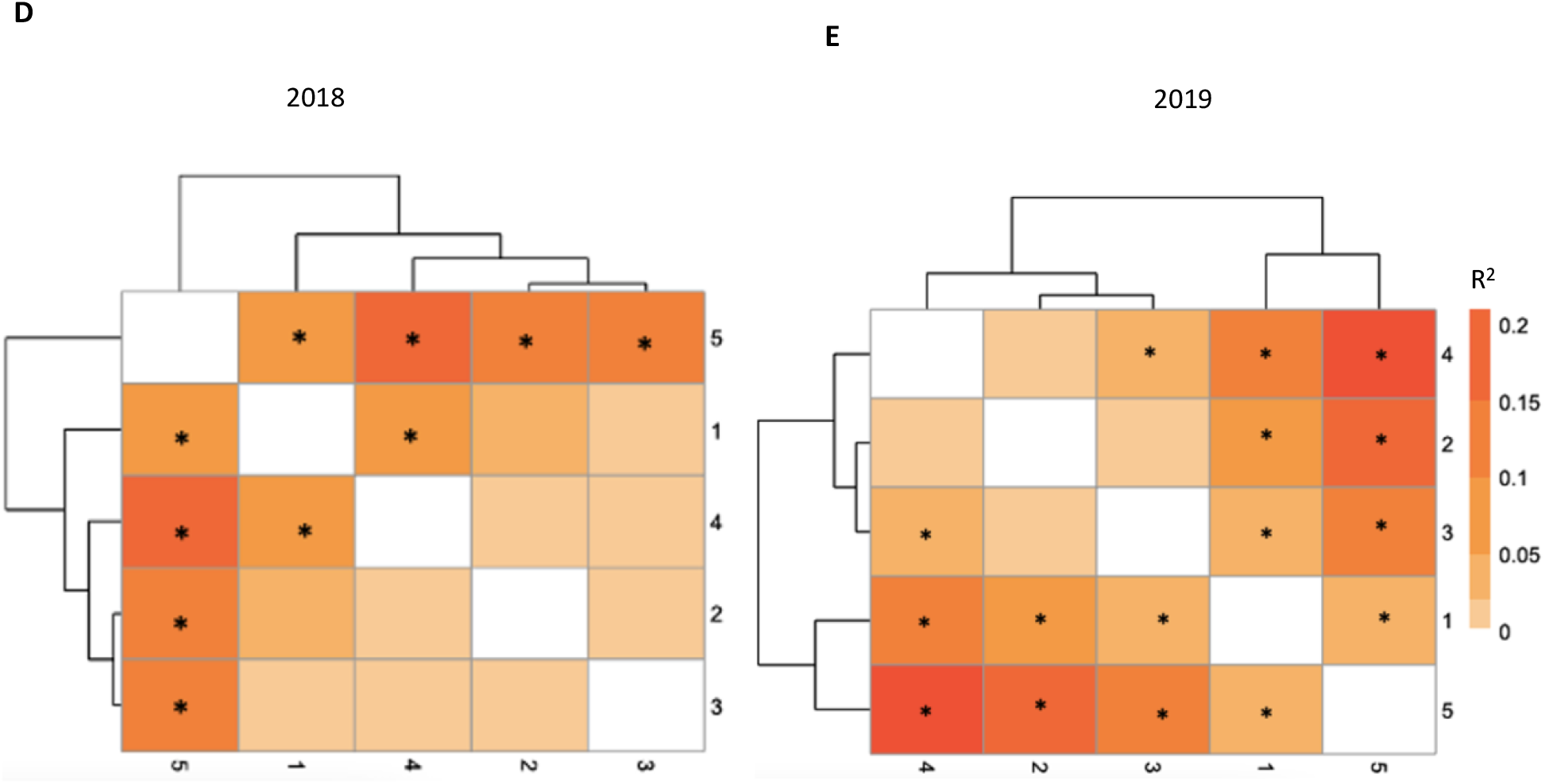
nMDS of Symbiodiniaceae types in *M. capitata* per regional block in Kāne‘ohe Bay in (A) 2018 and (B) 2019 and (C) comparing both years. Heatmap of the R^2^ of the pairwise PERMANOVA of the Symbiodiniaceae diversity per block in 2018 (D) and in 2019 (E). Significant terms are marked with *

**Table 1.**
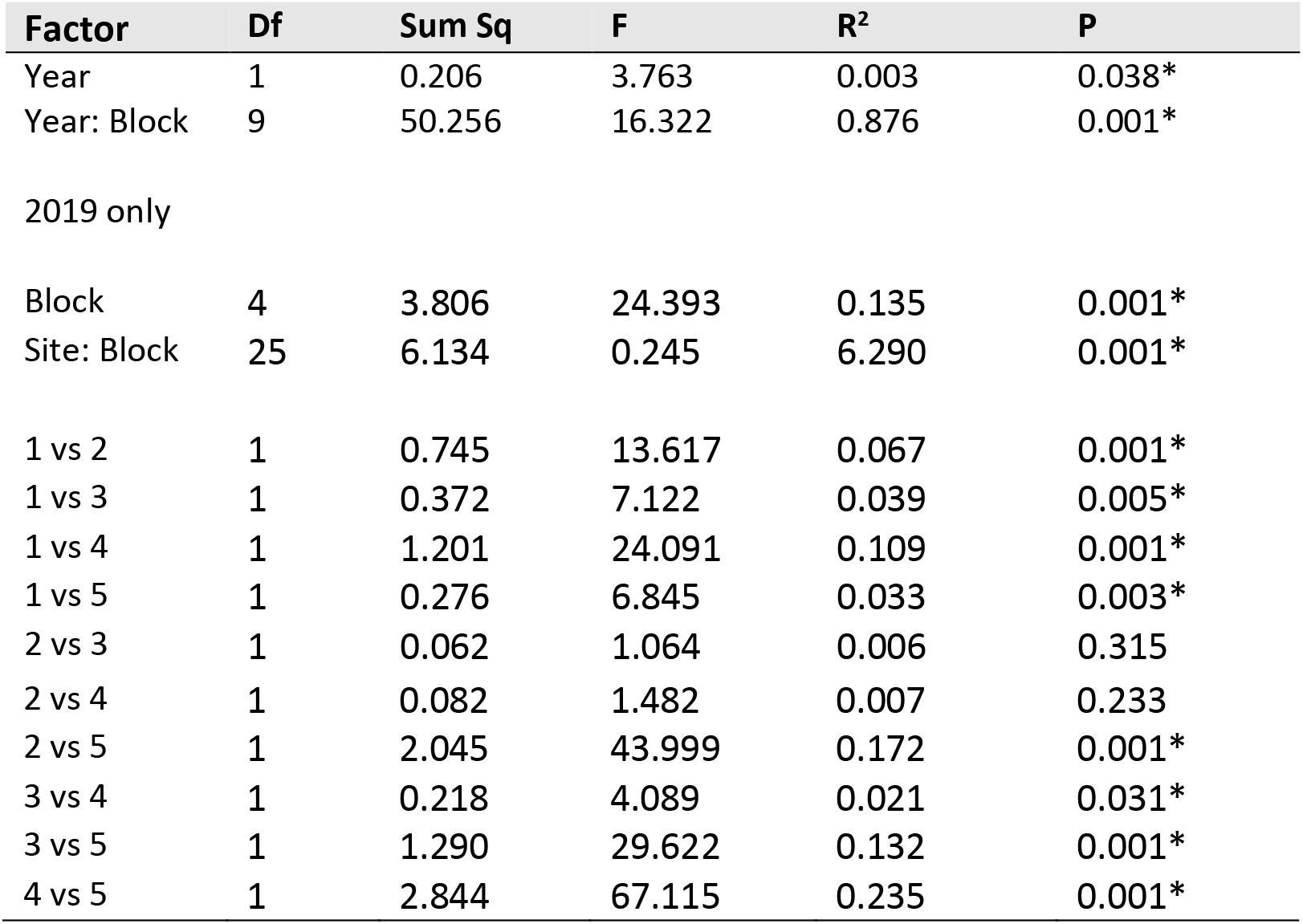
PERMANOVA based on Bray-Curtis dissimilarities of the *M. capitata* algal symbiont diversity present in corals sampled randomly from each environmentally defined block in Kāne‘ohe Bay, considering 2018 and 2019 years, and 2019 only. Permanova results for 2018 only are present in de Souza et al (81).

| Factor | Df | Sum Sq | F | $R^2$ | P |
| --- | --- | --- | --- | --- | --- |
| Year | 1 | 0.206 | 3.763 | 0.003 | 0.038* |
| Year: Block | 9 | 50.256 | 16.322 | 0.876 | 0.001* |
| 2019 only |  |  |  |  |  |
| Block | 4 | 3.806 | 24.393 | 0.135 | 0.001* |
| Site: Block | 25 | 6.134 | 0.245 | 6.290 | 0.001* |
| 1 vs 2 | 1 | 0.745 | 13.617 | 0.067 | 0.001* |
| 1 vs 3 | 1 | 0.372 | 7.122 | 0.039 | 0.005* |
| 1 vs 4 | 1 | 1.201 | 24.091 | 0.109 | 0.001* |
| 1 vs 5 | 1 | 0.276 | 6.845 | 0.033 | 0.003* |
| 2 vs 3 | 1 | 0.062 | 1.064 | 0.006 | 0.315 |
| 2 vs 4 | 1 | 0.082 | 1.482 | 0.007 | 0.233 |
| 2 vs 5 | 1 | 2.045 | 43.999 | 0.172 | 0.001* |
| 3 vs 4 | 1 | 0.218 | 4.089 | 0.021 | 0.031* |
| 3 vs 5 | 1 | 1.290 | 29.622 | 0.132 | 0.001* |
| 4 vs 5 | 1 | 2.844 | 67.115 | 0.235 | 0.001* |

### 3.3. SYMBIODINIACEAE SPATIAL VARIATION

There were significant differences in Symbiodiniaceae community composition among blocks of Kāne‘ohe Bay observed after thermal stress in 2019 (PERMANOVA, F _25_= 6.290, *p=* 0.001; **Table 1**). Although the large sample size of this study enables the detection of significant differences in overall community between years, the pairwise patterns between blocks remain consistent with results from 2018 (81). Blocks 1 and 5 were significantly different (**Figure 4D**, **Table 1**) from each other and the center region of the bay, while blocks in the middle of the bay were largely indistinguishable (except for block 3 vs 4, **Table 1**). Compared to 2018, bleaching in 2019 intensified the differences among these two spatial groups (81).

Despite a significant increase in the proportional representation of *Durusdinium* among the algal symbiont communities following bleaching, the underlying signal of geographic structure remained (**Figure 4A, B**).

### 3.4 DRIVERS OF SYMBIONT COMMUNITY COMPOSITION

The 30 sites had broadly different environmental characteristics. Depth varied from 0.5 m to 3.5 m; block 5 was the deepest block (mean 2.71 m), while sites in block 2 were shallowest (mean 1.36 m; Supplemental material **Table 1, 2**). Sedimentation ranged nearly 300-fold from 0.01 g/day to 2.93 g/day. In both years, sites in the middle of the bay had a smaller daily temperature range and daily temperature standard deviation when compared to sites in the northern and southern extremes of the bay (1 and 5). In 2019, block 4 had the highest increase in mean temperature and had the maximum absolute temperature (Supplemental material **Table 1, 2**).

**Table 2.** PERMANOVA of the environmental drivers of Bray-Curtis dissimilarities among Symbiodiniaceae communities in *M. capitata* among Kāne‘ohe Bay sites. Relative contribution was calculated as the sum square of each environmental factor divided by the sum of all environmental sum squares. Environmental factors explain 21% of Symbiodiniaceae variation.

| Environmental factors | Df | Sum Sq | F | P | Relative contribution |
| --- | --- | --- | --- | --- | --- |
| *Depth (m)* |  |  |  |  |  |
| Nominal | 1 | 17.585 | 156.348 | 0.001* | 0.6246 |
| *Temperature (°C)* |  |  |  |  |  |
| DHW | 1 | 0.382 | 3,967 | 0.040* | 0.0329 |
| Mean | 1 | 0.171 | 1.520 | 0.211 | 0.0073 |
| Maximum | 1 | 0.164 | 1.459 | 0.197 | 0.0104 |
| Minimum | 1 | 1.475 | 13.110 | 0.001* | 0.0116 |
| Daily range | 1 | 0.160 | 1.425 | 0.202 | 0.0205 |
| Mean daily standard deviation | 1 | 6.339 | 56.358 | 0.001* | 0.1889 |
| *Sedimentation (g/day)* |  |  |  |  |  |
| Mean | 1 | 0.059 | 0.522 | 0.553 | 0.0019 |
| Maximum | 1 | 0.139 | 1.231 | 0.267 | 0.0049 |
| Minimum | 1 | 0.239 | 2.123 | 0.126 | 0.0419 |
| Mean daily standard deviation | 1 | 1.008 | 8.958 | 0.001* | 0.0006 |
| Total |  | 27.721 |  |  |  |
| Residual |  | 99.878 |  |  |  |

We used dbRDA to examine environmental drivers of Symbiodiniaceae community structure and found six factors were significant after multiple comparisons correction (p<0.05; **Figure 5, Table 2**). In order of decreasing variance explained, depth, mean daily standard deviation in temperature, minimum temperature, sedimentation standard deviation, degree heating weeks significantly impacted Symbiodiniaceae community. Interestingly, most of these factors, were determined to be the major environmental drivers of Symbiodiniaceae community composition prior to the bleaching event (81).

**Figure 5.**
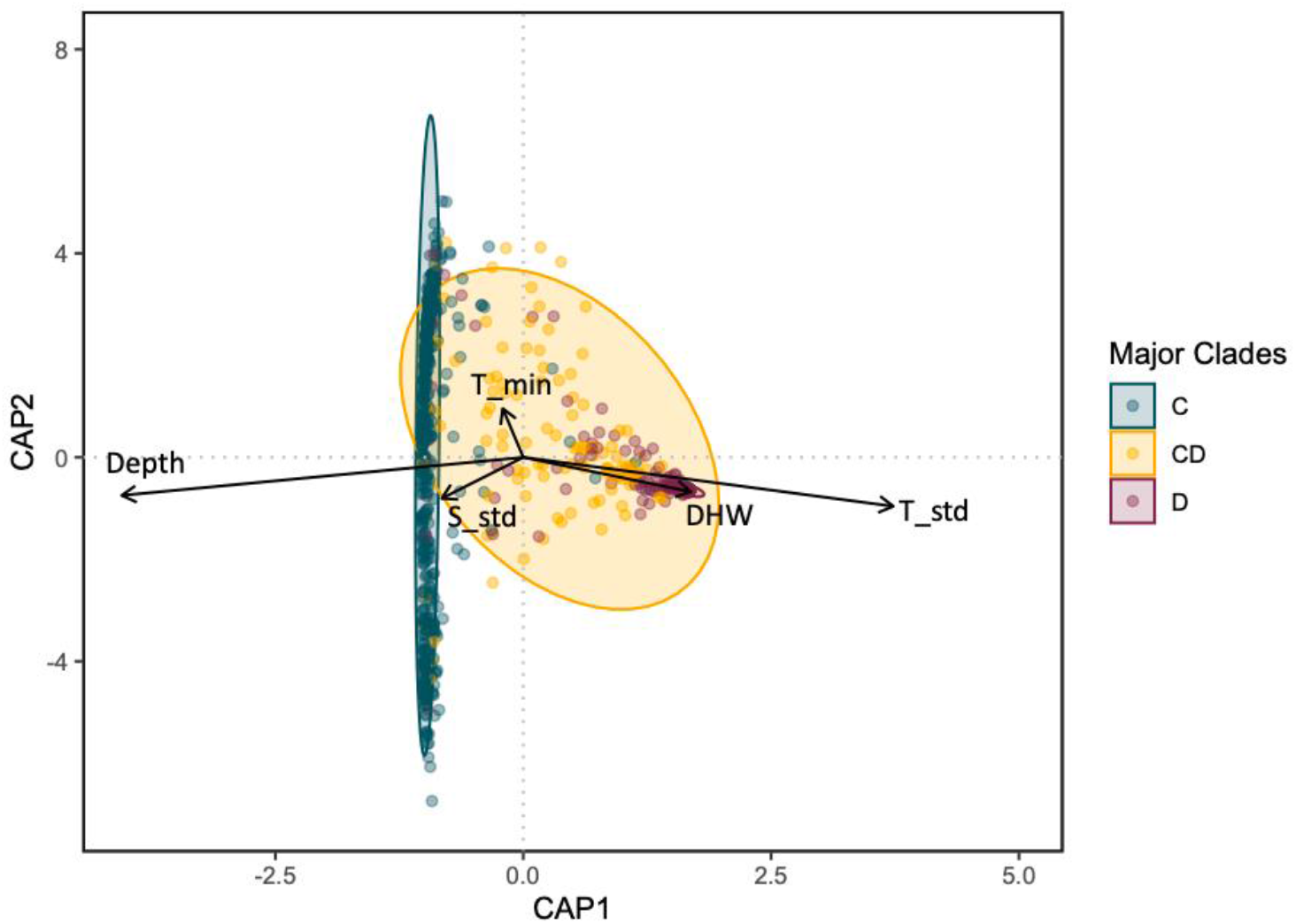
Distance based redundancy analysis (dbRDA) for environmental drivers of the Symbiodiniaceae communities measured in *Montipora capitata* in Kāne‘ohe Bay after the 2019 bleaching event. Each point represents a *M. capitata* colony sampled irrespective of site. For visualization, samples were considered as majority *Cladocopium* (C) if they contain > 80%C, majority *Durusdinium* (D) if > 80% D, and mixed CD otherwise. Only vectors for the environmental factors contributing significantly to the algal symbiont diversity are plotted. Each arrow signifies the multiple partial correlation of the environmental driver in the RDA whose length and direction can be interpreted as indicative of its contribution to the explained variation. T_std (temperature daily standard deviation), DHW (degree heating weeks), T_min (minimum temperature), DHW (degree heating weeks), S_std (sedimentation standard deviation).

## 4 Discussion

Temperature and irradiance are generally the main environmental factors underlying breakdown in the symbiosis between the coral host and their endosymbiotic community which drives coral bleaching (18, 54–57). However, intraspecific variability in bleaching severity within and among sites is also well documented (41, 44, 58–60), including in our study system (20, 41, 52). The extent to which endosymbiotic Symbiodiniaceae community composition contributes to such bleaching variability remains uncertain: some corals maintain stable associations throughout bleaching or revert to pre-bleaching algal symbiont composition (29, 30), whereas others maintain an altered algal symbiont community composition following bleaching events (31, 32). While there has been considerable research examining coral algal symbiont communities, the role of environmental factors in contributing to variation in both algal symbiont community structure and coral bleaching responses remains comparatively understudied (46, 50, 52, 102–104).

We previously quantified the algal symbiont community of *M. capitata* in relation to environmental gradients throughout Kāne‘ohe Bay in 2018 (81). Here, we resampled the same colonies following a natural bleaching event to examine Symbiodiniaceae diversity across an environmental mosaic and evaluate the relative importance of acute and chronic environmental conditions on symbiont communities. Like previous studies that often report an increase in heat-tolerant algal symbiont lineages when exposed to stressful conditions (34, 77, 105–108), we found a significant increase in *Durusdinium* when comparing Symbiodiniaceae communities of individually marked *M. capitata* colonies sampled before (early 2018) and shortly after (October 2019) the bleaching event. In addition, we show that the proportion of *Durusdinium* in a coral is negatively correlated with bleaching severity, supporting the potential for positive fitness consequences after bleaching if corals acquire thermally tolerant symbionts, consistent with the adaptive bleaching hypothesis (109). Cunning et al (110) found that at intermediate or low stress, corals decrease their proportion of heat stress algal symbionts (*Durusdinium*), while at higher stress (severe bleaching), the proportion of *Durusdinium* increases. Similarly, we found that block 4 had the highest maximum temperature increase, the most bleaching, and showed the greatest increase in *Durusdinium* compared to colonies in the other blocks.

Consistent with our previous study, the coral algal symbiont association was strongly influenced by environmental gradients. *Montipora capitata* located at the extreme southern and northern portions of Kāne‘ohe Bay hosted Symbiodiniaceae communities that were significantly different from the center of the bay and may be reflective of the unique environments in these regions. Despite a significant increase in the overall proportion of *Durusdinium* following the coral bleaching event, the same factors (depth, variability in temperature and sedimentation) emerge as significant drivers of Symbiodiniaceae community composition. In fact, 2018 and 2019 occupy almost the same multidimensional scaling space regardless of the bleaching event. Corals dominated by *Durusdinium* are much more similar (closer together the distance-based redundancy analysis) than those dominated by *Cladocopium* or with a mixed algal symbiont community composition, which corresponds to the reduced diversity of *Durusdinium* types and profiles we observed. Overall, these results suggest that the algal symbiont community composition was altered due to bleaching stress, but that such change was relatively minor in comparison to the established differences in community structure observed across the consistent environmental gradient of Kāne‘ohe Bay.

Our result is concordant with Dilworth et al. (42), who found consistency among the sites in the relative proportion of *Cladocopium* and *Durusdinium* in Kāne‘ohe Bay. Interestingly, Dilworth et al. (42), following a short thermal stress, in the mixed colonies, report a loss of *Cladocopium*, and higher recovery of *Durusdinium* compared to *Cladocopium* in bleached colonies. This finding may suggest that the algal symbiont community composition was starting to change but was not significant within the short time frame of the experiment. Alternatively, not all colonies may show such changes and this variability among individuals could make it difficult to detect significant changes. Cunning et al. (41) surveyed 60 colonies of *M. capitata* 6 months after the 2014 mass bleaching event and found that the dominant algal symbiont genus remained the same for 80% of colonies throughout their study. The remaining 20% showed variability in the dominant algal symbiont genus through time, but the changes that occurred were in either direction (i.e., both C to D and D to C) and were not related to visual bleaching. However, with relatively few colonies sampled, and less than 20% showing a change in dominant algal symbiont type, there is relatively little power in these previous studies to determine significance of directionality. In the hundreds of colonies sampled here, we show a slight but significant overall increase in the proportion of *Durusdinium* following the bleaching event. Consistent with these previous studies, the identity of the coral host and the local environmental history both appear to be important drivers of algal symbiont community composition, because despite a slight general increase in the proportion of *Durusdinium*, there remains a strong and essentially unchanged signature of the original environmental gradient on algal symbiont community from prior to the bleaching event (81). *M. capitata* is a vertical transmitter that releases symbiont provisioned eggs (111), creating a tight co-evolutionary linkage between host and symbionts. Host genetic differentiation in functional ontologies is also strongly associated with symbiont community (92), creating the potential for an environmental-symbiont linkage mediated by local adaptation of the host or holobiont.

While the increase in temperature during 2019 is an acute stress that led to bleaching, corals in different parts of the bay are also exposed to long-term environmental conditions that can act as chronic stressors, and might explain the mosaic spatial pattern of bleaching and Symbiodiniaceae composition across Kāne‘ohe Bay. For example, in other studies, corals with a long history of exposure to variable microhabitats were more heat tolerant than nearby conspecifics sampled from more stable regimes (36, 112–117). This resilience imparted by more variable environments may result from an “ecological memory” (118, 119) that can play a significant role in determining how individual corals will respond to a given stress. Environmental memory in corals following consecutive events has been documented in a few studies (e.g. 82, 121, 122). Although most mechanisms of environmental memory may be driven by the coral host (51), a change in algal symbiont composition (either shuffling of relative proportions or shifting to novel symbionts) may also play an important role. However, the relevance of such environmental memory remains controversial, with some studies suggesting a short duration, as corals revert to their initial algal symbiont compositions (108), whereas others report transgenerational inheritance of shuffled symbionts and suggest such change has major ecological relevance (122). It is interesting to note that corals in site 5_3 and 5_6 in block 5 did not host *Durusdinium* in any of the years. Possible explanations for this pattern include 1) *Durusdinium* is not available for the corals in those sites, 2) environmental drivers favor the selection of *Cladocopium* versus *Durusdinium* in those sites, or 3) the algal symbiont composition in those locations is driven by the host genetics, where local adaptation creates a tradeoff to hosting this genus. While Caruso et al (86) did not find a pattern of clonality in Kāne‘ohe Bay, when surveying the same colonies, it is unclear what genetic signals in the host would drive the Symbiodiniaceae composition. Taken together with our findings here, these studies indicate that Symbiodiniacae community composition responds to some combination of acute and chronic stressors, and that a better understanding of differences among host and algal symbiont species as well as drivers of environmental memory will improve our ability to predict coral bleaching at the level of individual colonies.

### Conclusions

Resampling of individually marked *M. capitata* colonies across the environmental mosaic of Kāne‘ohe Bay following a natural bleaching event revealed that patterns of algal symbiont distribution change as predicted. A marine heatwave resulted in a significant increase in the proportion of *Durusdinium* detected overall. However, there is strikingly little change in either the primary environmental drivers of Symbiodiniaceae community structure, nor the relative magnitude of those drivers in a distance-based redundancy analysis following the bleaching. The primary drivers of Symbiodiniaceae community structure remained consistently associated with depth and daily temperature variability despite the increased representation by *Durusdinium* in response to a bleaching event. This consistency in algal symbiont communities across chronic environmental gradients suggests that while communities may respond to short-term acute stressors such as heat waves, they appear constrained by the long-term environmental conditions surrounding the holobiont.

5

## Acknowledgments

We thank the Coral Resilience Lab (the legacy of Ruth Gates) and ToBo Lab for assistance with collections, molecular work or bioinformatics (special thanks to Kira Hughes, Joshua Hancock, Janaya Bruce, Dennis Conneta, Caroline Hobbs, Valerie Kahkejian, and Christian Marin). This work was funded by a Coordenação de Aperfeiçoamento de Pessoal de Nível Superior - Brasil (CAPES) fellowship, the National Science Foundation (OA-1416889 & BioOCE-1924604 to RJT) and the Paul G. Allen Family Foundation.

## Compliance with ethical standards

Conflict of interest: On behalf of all authors, the corresponding author states that there is no conflict of interest.

## Contributions

M.R.S, C.C., L.R.J., R.G. conceived of and designed the experiment, and M.R.S, C.C. & L.R.J. led the collections. M.R.S, C.C., L.R.J. contributed to molecular work. M.R.S., C.D., R.J.T. contributed to analysis; C.D., R.G., R.J.T. contributed to funding. The first draft of the paper was written by M.R.S with help from R.J.T. All authors contributed to discussion, data interpretation, manuscript revisions, and approved the final manuscript.

