## Supplemental material for "Coral environmental history is primary driver of algal symbiont composition, despite a mass bleaching event"

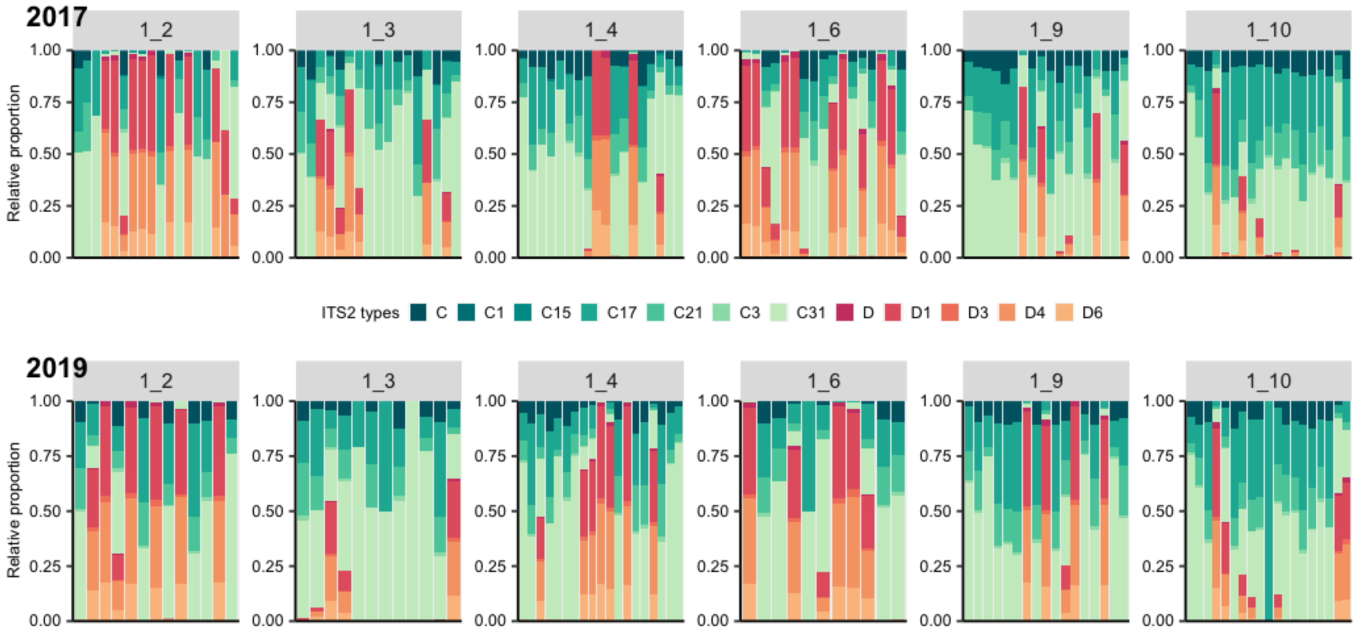


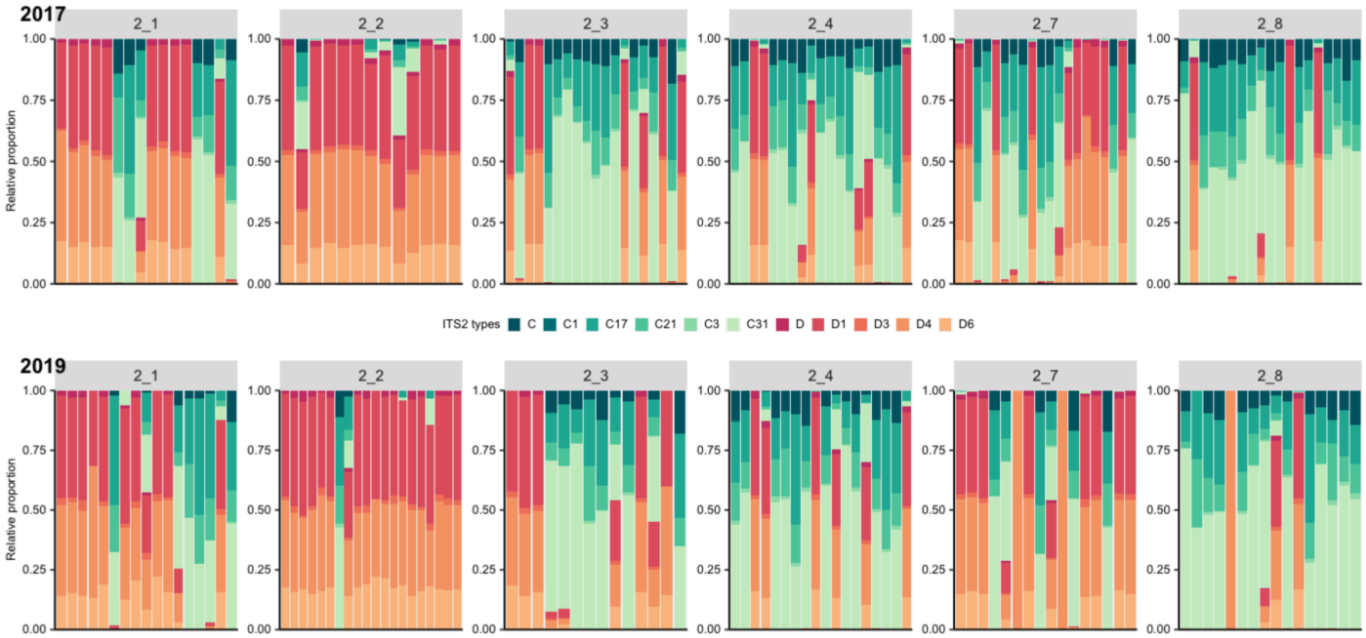

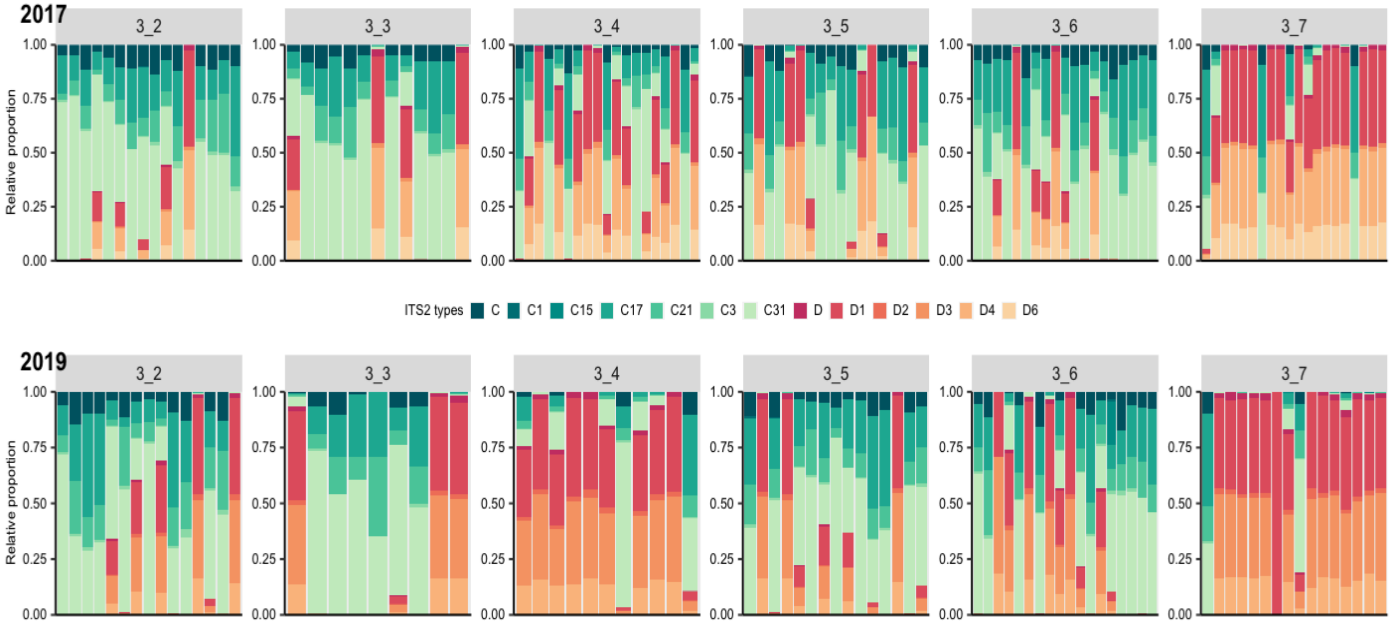
2017
3_2
3_3
3_4
3_5
3_6
3_7
Relative proportion
1.00
0.75
0.50
0.25
0.00
ITS2 types C C1 C15 C17 C21 C3 C31 D D1 D2 D3 D4 D6
2019
3_2
3_3
3_4
3_5
3_6
3_7
Relative proportion

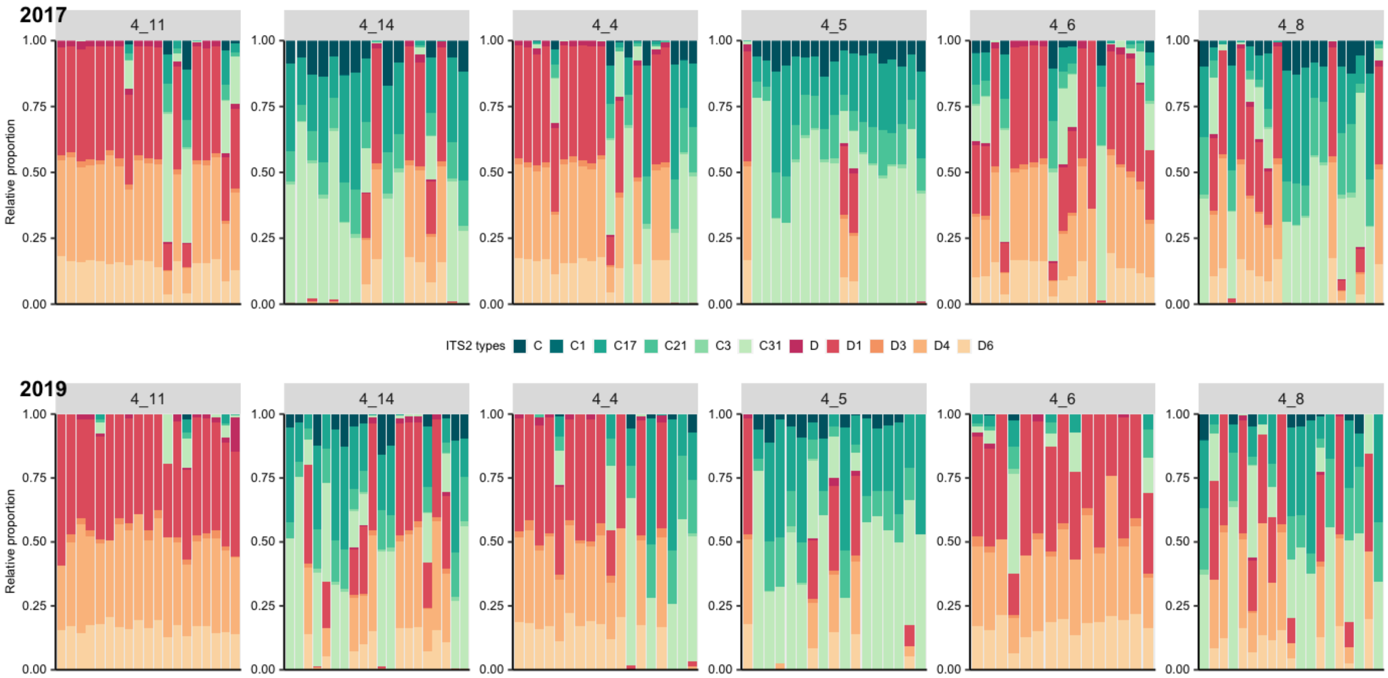
2017
4_11
4_14
4_4
4_5
4_6
4_8
Relative proportion
1.00
0.75
0.50
0.25
0.00
ITS2 types C C1 C17 C21 C3 C31 D D1 D3 D4 D6
2019
4_11
4_14
4_4
4_5
4_6
4_8
Relative proportion

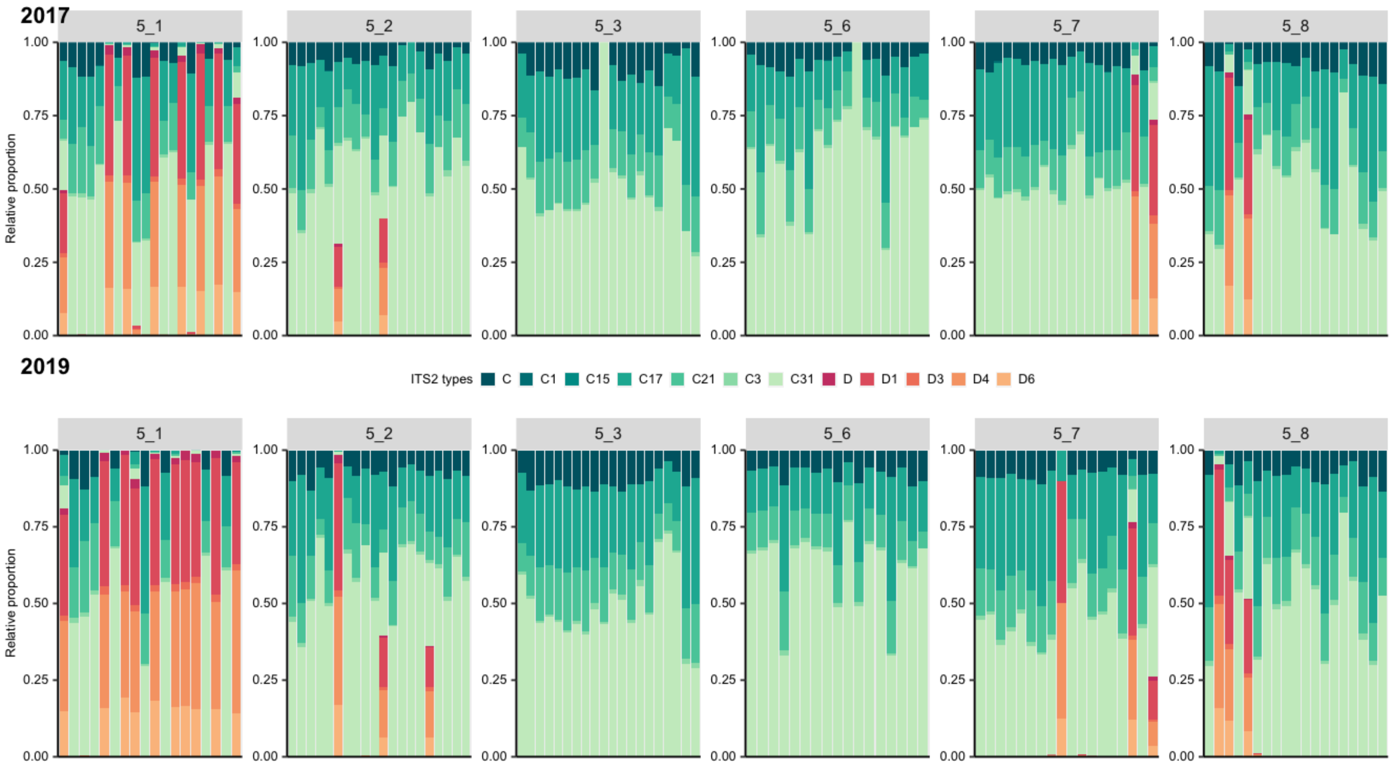


**Figure 1.** Relative proportion of Symbiodiniaceae types present in each *Montipora capitata* colony in each site. Panels shows the different blocks - block 1 (A), block 2 (B), block 3 (C), block 4 (D), block 5 (E).

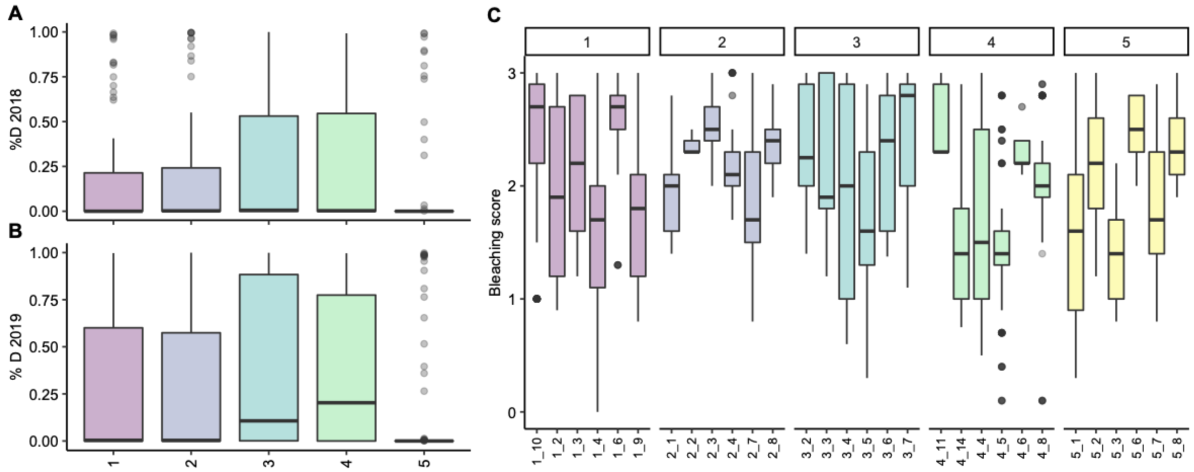


**Figure 2.** Proportion of *Durusdinium* per block in A) 2018 and B) 2019. C) Bleaching score per site.

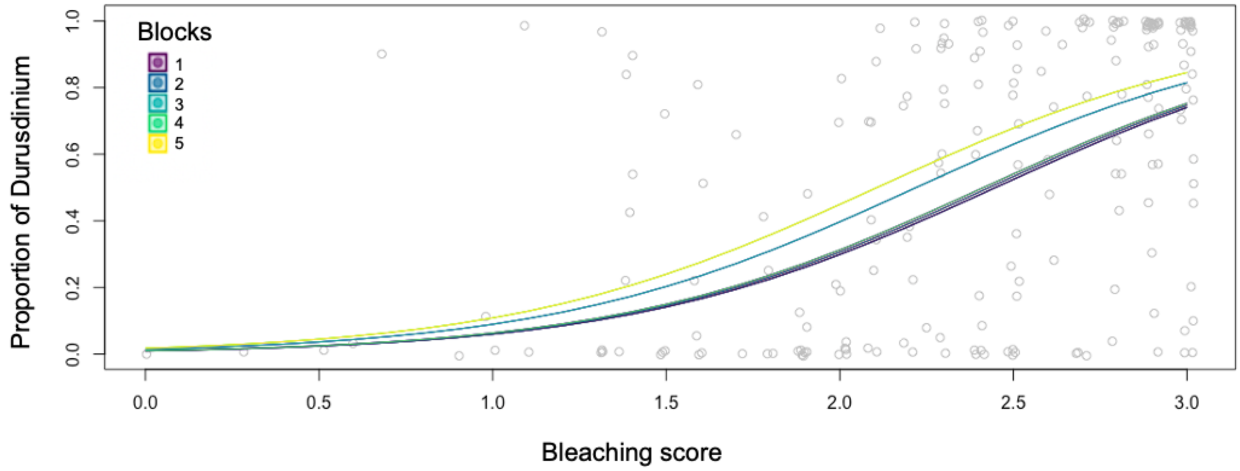


**Figure 3.** General linear regression of the proportion of *Durusdinium* per bleaching score, colored per block.

**Table 1**. Site information, sedimentation, and comparison of temperature for 2018 and 2019.

| Site | Block | Depth (m) | Sedimentation mean g/day | Temperature max 2018 | Temp max 2019 |
|---|---|---|---|---|---|
| 1_2 | 1 | 1.2 | 0.06602 | 29.3728 | 29.7687 |
| 1_3 | 1 | 3 | 0.02187 | 28.8250 | 28.6231 |
| 1_4 | 1 | 3.5 | 0.01931 | 28.8416 | 29.7490 |
| 1_6 | 1 | 0.7 | 0.02287 | 28.9421 | 30.2274 |
| 1_9 | 1 | 2.6 | 0.02982 | 28.8125 | 28.7719 |
| 1_10 | 1 | 3 | 0.02928 | 28.8859 | 28.6366 |
| 2_1 | 2 | 1.2 | 0.16410 | 28.8539 | 29.4408 |
| 2_2 | 2 | 0.5 | 0.04501 | 28.8794 | 28.7067 |
| 2_3 | 2 | 0.5 | 0.02779 | 29.2753 | 30.1186 |
| 2_4 | 2 | 2.4 | 2.93056 | 28.6209 | 29.0228 |
| 2_7 | 2 | 1.5 | 0.40130 | 28.7231 | 30.5101 |
| 2_8 | 2 | 2.1 | 0.27611 | 28.7231 | 28.9946 |
| 3_2 | 3 | 1.4 | 0.09804 | 28.1498 | 28.9693 |
| 3_3 | 3 | 1 | 0.05737 | 28.9506 | 28.6542 |

| | | | | | |
|---|---|---|---|---|---|
| 3_4 | 3 | 1 | 0.04163 | 29.2845 | 29.4586 |
| 3_5 | 3 | 0.9 | 0.03061 | 29.0320 | 30.1222 |
| 3_6 | 3 | 0.9 | 0.03796 | 28.9826 | 31.9997 |
| 3_7 | 3 | 0.7 | 0.02244 | 29.1680 | 29.6930 |
| 4_4 | 4 | 1.4 | 0.04811 | 28.9492 | 30.0067 |
| 4_5 | 4 | 1.1 | 0.03305 | 28.9298 | 29.6440 |
| 4_6 | 4 | 1.5 | 0.34257 | 29.1865 | 31.1699 |
| 4_8 | 4 | 1.2 | 0.01573 | 28.8363 | 29.9775 |
| 4_11 | 4 | 1.4 | 0.07752 | 29.2206 | 30.1610 |
| 4_14 | 4 | 1.4 | 0.16850 | 28.9260 | 30.0172 |
| 5_1 | 5 | 1.2 | 0.21580 | 29.3822 | 29.7978 |
| 5_2 | 5 | 2.7 | 0.62898 | 28.8171 | 28.5985 |
| 5_3 | 5 | 2.4 | 0.13268 | 28.7982 | 28.5721 |
| 5_6 | 5 | 3.5 | 0.41842 | 27.6119 | 30.0287 |
| 5_7 | 5 | 3.3 | 0.38405 | 28.6961 | 29.6721 |
| 5_8 | 5 | 3.2 | 0.28655 | 28.7486 | 29.6730 |

**Table 2.** Summary statistics per block in 2018 and 2019

| Block | Mean °C 2018 | Mean °C 2019 | Max °C 2018 | Max °C 2019 | Daily °C range 2018 | Daily °C range 2019 | Daily °C standard deviation 2018 | Daily °C Standard deviation 2019 |
|---|---|---|---|---|---|---|---|---|
| **1** | 25.697 | 26.207 | 28.946 | 29.296 | 0.244 | 1.007 | 0.243 | 0.264 |
| **2** | 25.647 | 26.265 | 28.834 | 29.465 | 1.246 | 1.333 | 0.335 | 0.355 |
| **3** | 25.798 | 26.335 | 28.928 | 29.816 | 1.871 | 2.014 | 0.474 | 0.505 |
| **4** | 25.716 | 26.536 | 29.008 | 30.162 | 1.261 | 1.383 | 0.342 | 0.380 |
| **5** | 25.490 | 26.159 | 28.675 | 29.390 | 0.746 | 0.756 | 0.204 | 0.212 |
